# Visualizing and quantifying structural diversity around mobile resistance genes

**DOI:** 10.1101/2023.08.07.551646

**Authors:** Liam P. Shaw, Richard A. Neher

## Abstract

Understanding the evolution of mobile genes is important for understanding the spread of antimicrobial resistance (AMR). Many clinically important AMR genes have been mobilized by mobile genetic elements (MGEs) on the kilobase scale, such as integrons and transposons, which can integrate into both chromosomes and plasmids and lead to rapid spread of the gene through bacterial populations. Looking at the flanking regions of these mobile genes in diverse genomes can highlight common structures and reveal patterns of MGE spread. However, historically this has been a largely descriptive process, relying on gene annotation and expert knowledge. Here we describe a general method to visualize and quantify the structural diversity around genes using *pangraph* to find blocks of homologous sequence. We apply this method to a set of twelve clinically important beta-lactamase genes and provide interactive visualizations of their flanking regions at https://liampshaw.github.io/flanking-regions. We show that nucleotide-level variation in the mobile gene itself generally correlates with increased structural diversity in its flanking regions, demonstrating a relationship between rates of mutational evolution and rates of structural evolution, and find a bias for greater structural diversity upstream. Our framework is a starting point to investigate general rules that apply to the horizontal spread of new genes through bacterial populations.

**Impact statement:** Understanding the evolution and spread of mobile resistance genes is challenging because of the high variability in their genomic contexts. Here we outline a fast computational approach that identifies stretches of homologous sequence in the flanking regions of a gene, simultaneously producing interactive visualizations of these regions and quantifying the diversity within them. As an example, we apply the method to twelve clinically important beta-lactamase genes. We find that structural diversity around the resistance gene is correlated with mutations within it and that there is greater structural diversity upstream of many genes. There may be other general patterns about the evolution of mobile resistance genes that can be recovered with this kind of analysis.

## I. INTRODUCTION

Mobile genetic elements (MGEs) allow the horizontal movement of genes within and between bacterial species, contributing to a vast range of bacterial phenotypes including antimicrobial resistance (AMR). A ‘mobile gene’ will be found in multiple genomic contexts, where its flanking regions contain information on the evolutionary history of the MGE (or MGEs) that have mobilized it.^1^ However, despite the increasing availability of complete genomes, high structural diversity in these flanking regions generated by both transposition and recombination means that analysing them is challenging.

Each mobile AMR gene has its own unique epidemiological history. As a recent example, the metallobeta-lactamase gene *bla*_NDM−1_ was first reported by Yong et al. (2009a). The earliest NDM-positive isolate dates only from 2005 (Jones et al., 2014), but already by the 2010s *bla*_NDM−1_ had been seen worldwide in diverse bacteria. As of 2023 there are public genomes containing *bla*_NDM_ genes from seventeen bacterial genera (Alcock et al., 2023). Such a rapid horizontal and global spread with multiple rearrangements presents a challenge for genomic epidemiology. The flanking regions around *bla*_NDM−1_ show large structural diversity, particularly upstream of the gene, although with some traces of a common ancestral MGE. Acman et al. (2022) developed a methodology for iteratively ‘splitting’ flanking sequences, finding that downstream patterns of structural diversity in a global dataset supported the previous conclusion that *bla*_NDM−1_ had been first mobilized by a Tn*125* transposon (Toleman et al., 2012). This example demonstrates that analysis of flanking regions is a valuable but difficult task, often requiring bespoke methods. For this reason, almost all the existing literature on the flanking regions of mobile genes remains descriptive and focused on single genes at a time, meaning it is difficult to extrapolate the general rules – if any – that govern evolution in these regions.

The high levels of structural diversity around mobile genes present two challenges: visualization and quantification. Visualizations are often based on annotated gene clusters (Gilchrist and Chooi, 2021), but genes are frequently disrupted in flanking regions. Other tools aim to identify discrete clusters for tracking MGE epidemiology. For example, TETyper was developed specifically for transposable MGEs and can identify small-scale changes associated with transposition (Sheppard et al., 2018) and Flanker performs alignment-free clustering of flanking sequences based on mash distances (Matlock et al., 2021). Such tools are useful but difficult to connect to the processes that generate structural diversity. In general, our understanding of these processes remains qualitative.

Here, we aim to provide a starting point for quantitative analysis of structural diversity around mobile genes. We outline an annotation-free approach based on finding homologous sequence blocks with *pangraph* (Noll et al., 2023) that scales to thousands of sequences and connects visualization with quantification. As a demonstration, we apply our method to the flanking regions of twelve beta-lactamase genes.

**FIG. 1.**
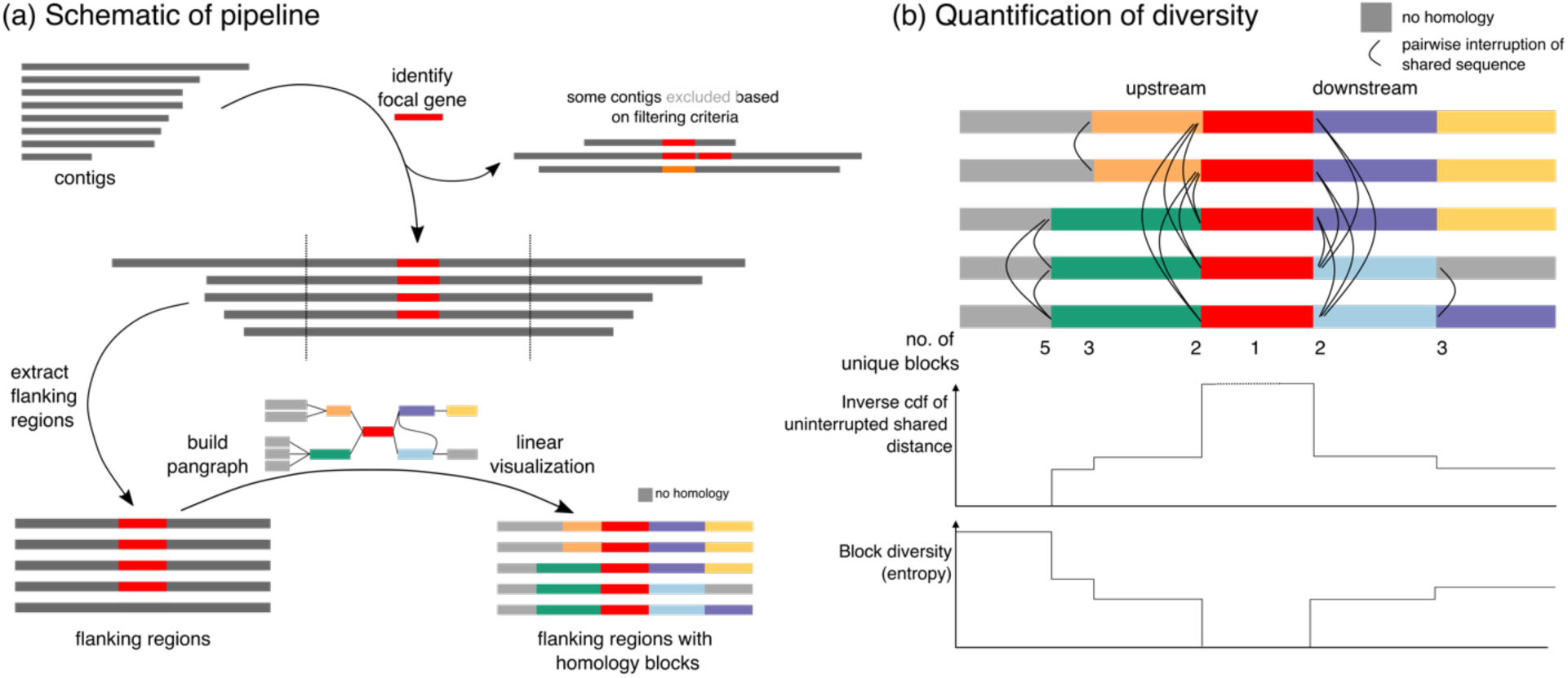
Overview of pipeline and quantification of structural diversity. (a) Given a focal gene and a set of contigs as input, the pipeline extracts the gene’s flanking regions (default: 5kb upstream/downstream). Contigs are filtered based on user-defined criteria, removing those with >1 copy of the gene, excessive diversity in the focal gene (default: >25 nucleotide-level differences), or inadequate lateral coverage of the gene (default: <99%). The extracted flanking regions are then passed into *pangraph* to identify blocks of homologous sequence and create a graph data structure. This graph data structure can be used to visualize the original flanking regions with blocks coloured by homology in a linear layout, highlighting shared homology across input contigs. Optionally, annotations (in gff format) can also be included. This linear visualization is produced as an interactive html output. Blocks without homology to other blocks in the input contigs are coloured grey (i.e. unique sequence). (b) For any pairwise comparison of flanking regions, we define the ‘uninterrupted shared distance’ as the upstream/downstream distance to their first breakpoint in homology i.e. indicated by different coloured blocks or by grey blocks. The inverse cumulative distribution function of these pairwise uninterruped shared distances shows how structural similarity falls away with distance from the focal gene in both directions. The block diversity, computed as the entropy of the vector of blocks at any linear position, gives another way of measuring structural diversity at distances from the focal gene.

## II. METHODOLOGY

### Overview of the pipeline

Figure 1a gives a schematic overview of the pipeline available at https://github.com/liampshaw/mobile-generegions. The use case is where an investigator has a *focal gene* and a set of *contigs*. For example, the focal gene might be *bla*_NDM−1_ and the contigs would be assemblies containing *bla*_NDM_ gene variants downloaded from NCBI MicroBIGG-E with the search query element_symbol:blaNDM*. The first step is to locate the focal gene, using blastn (default parameters) followed by retaining only hits matching lateral coverage and nucleotide-level difference thresholds which can be specified by the user (default: >99% coverage, <25 nucleotide-level differences in the alignment, with 1 single nucleotide variant counted as 1 difference and an *n*-bp indel counted as *n* differences; in practice, almost all variants are SNVs so we refer to this threshold as a SNV threshold). This location is then used to extract the flanking regions for some specified distance value (default: +/-5kb upstream and downstream). By default, the pipeline orientates the extracted region so that the focal gene is on the positive strand. It also omits contigs which contain more than one copy of the focal gene and those which are are shorter than the requested flanking distance (schematic examples of excluded contigs are shown in Figure 1a).

The pipeline then uses *pangraph* (Noll et al., 2023) to find homologous sequences. The *pangraph* algorithm was developed for whole genomes, and aims to identify stretches of homologous sequence within and across all input genomes, approximating multiple-genome alignment through iterative pairwise alignment of subsets of sequences with either *minimap2* or *mmseqs2*. These stretches of homologous sequence are referred to as ‘pancontigs’: linear multiple-sequence alignments, with small indels and nucleotide polymorphisms retained in the underlying data structure. The algorithm aligns pairs of subgraphs in a traversal of a guide tree of all input sequences, using a score to determine the most favorable mergers of aligned pancontigs which takes into account the alignment length, the creation of additional pancontigs, and the number of polymorphisms. The output of *pangraph* summarizes a set of input genomes into a highly condensed graph structure: pancontigs are connected by edges if adjacent in any input sequence, and individual genomes are represented by paths through the graph.

Here, we are applying *pangraph* to much shorter regions than whole genomes to arrive at a simplified representation of structural diversity in flanking regions, which are typically highly diverse and have many breaks in homology due to large-scale insertions and rearrangements. We refer to pancontigs as ‘homologous blocks’ of sequence and colour blocks according to this homology, using this to produce visualizations that show their coarse-grained structural arrangement. Because *pangraph* was developed with the Mb-scale of whole genomes in mind, it can find homologous blocks within kbs of diverse flanking regions across thousands of sequences on a personal computer in minutes: for a dataset of *n* = 1,581 *bla*_TEM_-positive assemblies from 24 bacterial genera, building the pangraph of ∼10kb flanking regions (+/5kb) takes less than 3 minutes on a laptop (2 GHz Quad-Core Intel Core i5 processors, 16 GB RAM).

**TABLE I.**
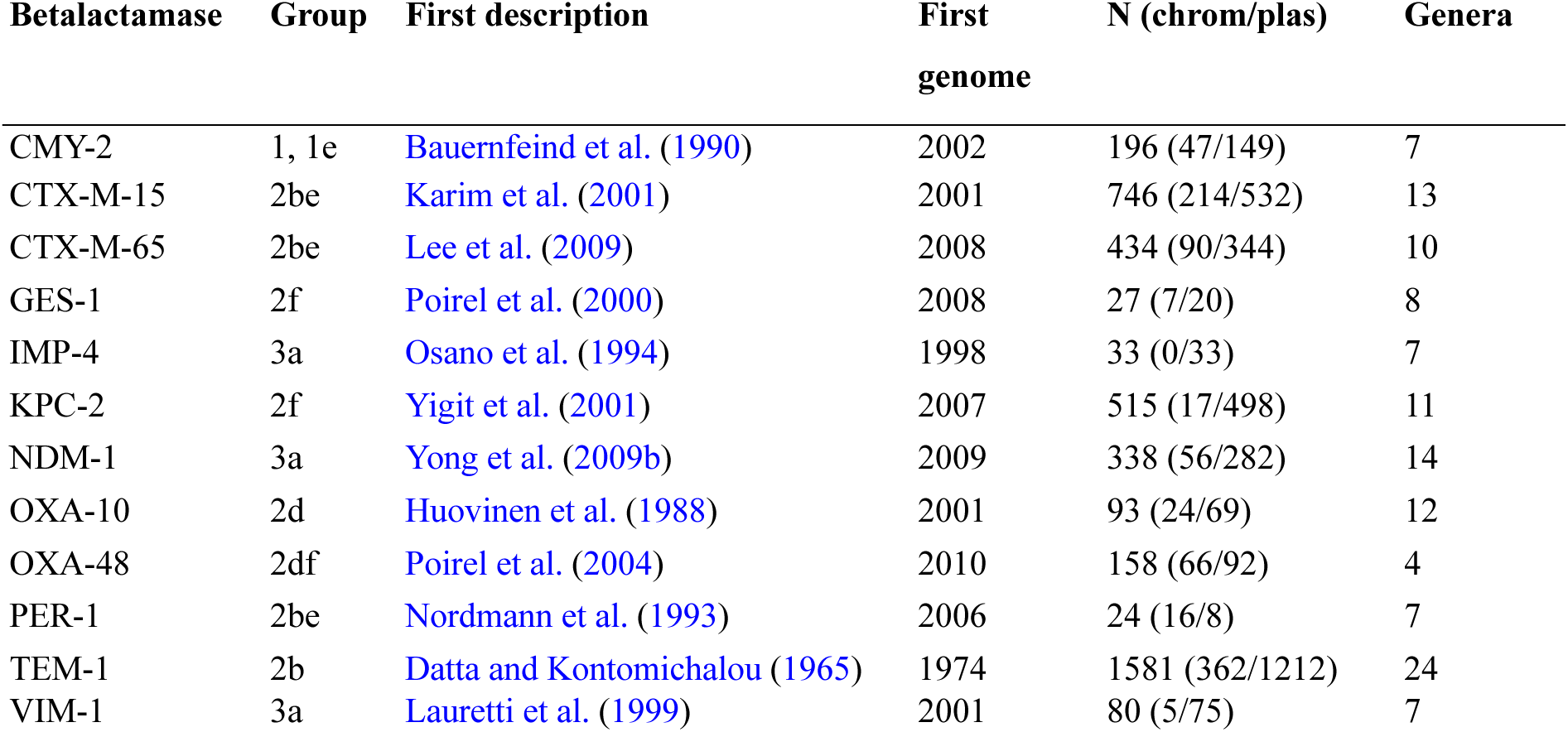
Beta-lactamase genes used as focal genes for example analysis. We started from the CARD prevalence database v3.1.0 and re-analysed sequences to confirm gene presence. Chromosome/plasmid designation is taken from CARD (‘ncbi_chromosome’ or ‘ncbi_plasmid’). Only those with sufficient flanking sequence (+/-5kb either side of the focal gene) with <25 nucleotide-level differences in the focal gene were taken forward for further analysis. Almost all variants were SNVs: of 6,186 gene sequences extracted across the dataset, only 87 (1.5%) were not exactly the same length as the focal gene. Functional group information is as in Bush and Jacoby (2010). First descriptions are given for the specific named focal gene rather than the enzyme family as a whole – for example, the CTX-M family was first described in 1990 (Bauernfeind et al., 1990).

Since the aim is to provide a coarse-grained representation of homology for visualisation and quantification, by default the pipeline uses a minimum block size of 100 bp with the *asm10* sensitivity level in *minimap2*. In the *pangraph* representation a block has a consensus sequence and thus a consensus length, though individual sequences may have insertions or deletions making them vary in length. This consensus sequence is used to compare and align blocks in the *pangraph* algorithm, which makes *pangraph* fast but may introduce minor inconsistencies and artifacts in the block alignments. Although we use the true lengths of sequences in all visualizations (meaning that identically coloured blocks may appear as slightly different lengths in visualizations), we do not analyse small-scale structural differences below the length scale of blocks. We note that it is possible to use this information for investigating structural rearrangement, such as using target site duplications (<15bp direct repeats) to indicate newly transposed copies of transposable elements as in TETyper (Sheppard et al., 2018), but we do not investigate them here. However, no sequence information is lost and a true multiple sequence alignment for every block can be reconstructed with *pangraph* using the ‘polish’ command for more detailed analysis (see *pangraph* documentation).

### Visualization

The pangraph of the flanking regions in GFA format can be viewed in Bandage, although for most mobile genes the default force-directed layout will produce an uninformative ‘hairball’ due to repeated homology blocks. We therefore produce a linear interactive html representation of the flanking region with blocks coloured by homology (Fig. 1a). Unique stretches of sequence with no homology within the dataset are coloured grey. Users can highlight a particular block by clicking on it. Optionally, if annotations are provided for the input sequences in gff format these can be projected onto this visualization and toggled on and off (for examples, see the beta-lactamase link).

### Quantification

The coarse-grained representation of the flanking regions in terms of homologous blocks is our starting point for quantitative analysis. We define two quantities, illustrated schematically in Fig. 1b:

- *Uninterrupted shared distance*: for a pairwise comparison of two flanking regions, the distance upstream/downstream from the focal gene from the focal gene to their first structural difference, i.e. the first point where their paths in the graph diverge. The inverse cumulative distribution function (cdf) of all pairwise uninterrupted shared distances in a set of sequences represents the decay of structural similarity with distance from the focal gene, and can help to identify common positions where non-homologous structural variation is introduced.
- *Block diversity*: at a given distance away from the focal gene, the Shannon entropy of the homologous blocks at that location across the dataset. If *p_i_*(*x*) is the proportion of sequences at distance *x* with block *i*, then 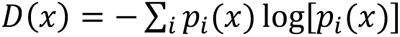.

### Beta-lactamase dataset

We first used prevalence information from the Comprehensive Antibiotic Resistance Database (CARD) v3.1.0 (Alcock et al., 2023) to assemble a dataset of genomic sequences that contained at least one gene encoding a beta-lactamase from any of twelve families (CMY, CTX-M, GES, IMP, KPC, NDM, OXA, PER, SHV, TEM, VIM, VEB) according to CARD’s ‘strict’ matching criteria and were coded by CARD as ‘ncbi_chromosome’ (n=3,199) or ‘ncbi_plasmid’ (n=4,026). We matched information from these NCBI accessions to get their BioSample ID and also used NCBI entrez to link them to metadata such as species, collection date, host, and geographical location (11 had no associated BioSample). We automatically assigned country names from the NCBI variable geo_loc_name and also from lat_lon where possible. We inspected 42 entries where automatic country names failed and inputted the country manually. Where samples were described as e.g. ‘USA ex Mexico’ we coded this as USA.

In the final cleaned metadata: 5,958 sequences (82.5%) had a collection year, 6,215 (86.0%) had a country, and 5,764 (79.8%) had both. At the level of taxonomic order, most sequences were from species within Enterobacterales (5,529/7,225, 76.6%). Beta-lactamase families can be diverse and contain non-homologous genes – notably, the OXA family. Therefore, we used a review of the literature and the classification of clinically important beta-lactamase families (Bush and Jacoby, 2010) to choose twelve clinically important beta-lactamase genes that have emerged recently as mobile AMR threats as focal genes for our example quantitative analysis (Table I). For these twelve focal genes, we included only sequences in our dataset that had sufficient flanking sequence either side of the focal gene (+/-5kb) with <25 nucleotide-level differences in the focal gene itself (n=3,362 total; sequences with length <10kb + gene length are omitted by this threshold e.g. small plasmids). This corresponds to a nucleotide identity cutoff ranging from 96.6% for *bla*_IMP−4_ (shortest gene, 741 bp) to 97.8% for *bla*_CMY−2_ (longest gene, 1,146 bp), although in practice nearly all included had <7 SNVs in the focal gene so were >99% identical at the nucleotide level.

We compiled this dataset in this way before we were aware of NCBI MicroBIGG-E. MicroBIGG-E allows users to query genomes already analysed with NCBI AMRFinderPlus as part of the Pathogen Detection Pipeline for a specific gene or genetic element, and then download only its flanking regions across all genomes. This makes it an ideal starting point for flanking region analysis with our pipeline. However, as of 16 November 2023 it only lets the user download flanking regions +/-2kb. Looking at larger flanking regions currently requires downloading the whole contigs first.

## III. APPLICATION: BETA-LACTAMASE GENES

To demonstrate the scalability of our method, we applied it to twelve different beta-lactamase genes (Table I). Beta-lactam antibiotics are key to modern medicine, accounting for 65% of prescriptions for injectable antibiotics in the United States (Bush and Bradford, 2016). These antibiotics share a common component: the beta-lactam ring, first seen in the structure of penicillin. Beta-lactamases are a diverse group of enzymes which can break apart the beta-lactam ring by hydrolysis and render beta-lactam antibiotics ineffective. The use of beta-lactam antibiotics therefore exerts a strong selective pressure for sensitive bacteria to carry betalactamases. This effect was first observed immediately after the widespread introduction of penicillin: at one London hospital the proportion of penicillin-resistant *Staphylococcus aureus* carrying the beta-lactamase *bla*Z increased from 14% in 1946 to 38% the following year (Barber, 1947). The increase in the prevalence of beta-lactamases and their adaptation to hydrolyse successive generations of beta-lactams to is one of the clearest real-world examples of rapid evolution.

**FIG. 2.**
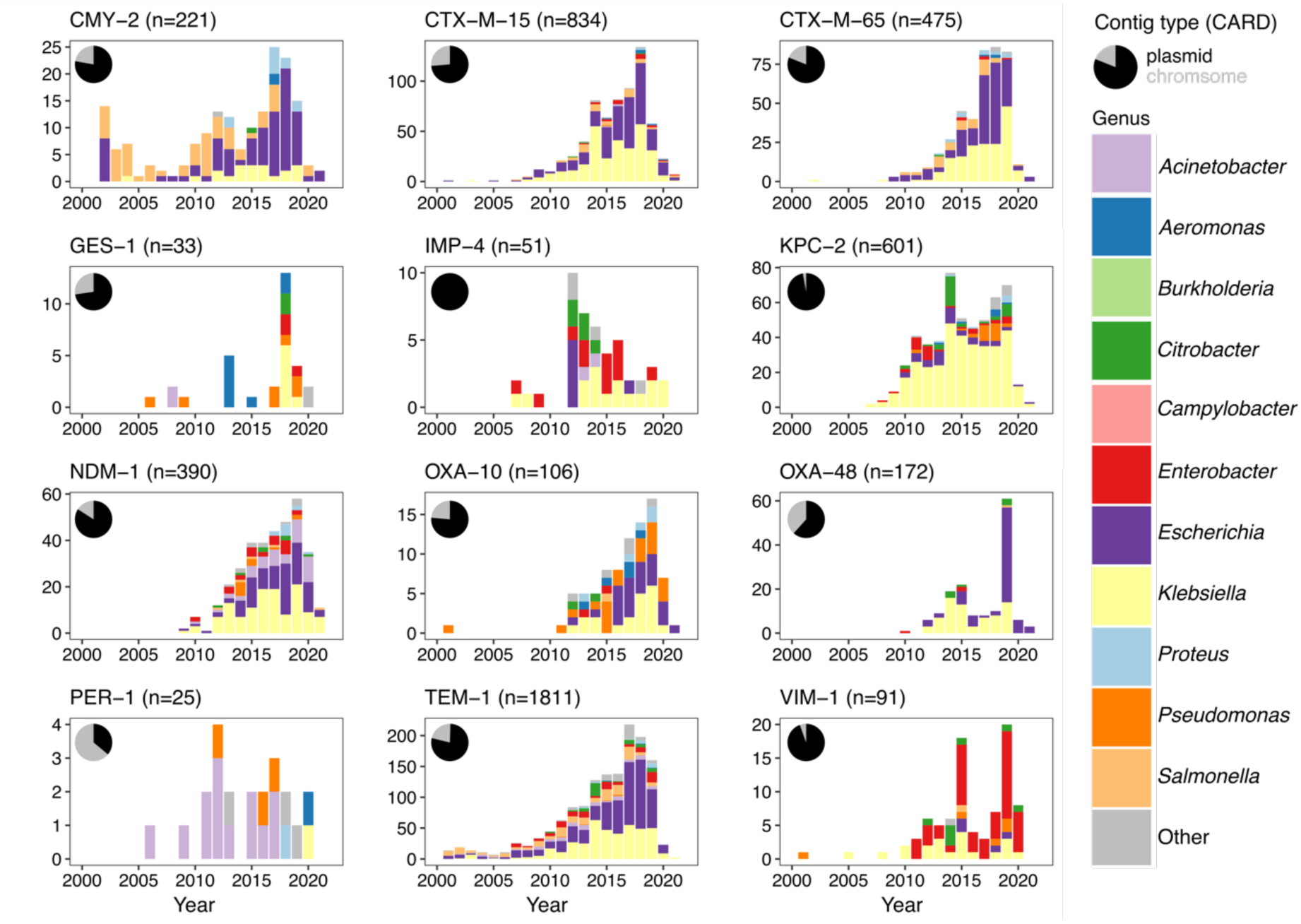
Summary of the flanking regions analysed for each beta-lactamase. Inset pie-charts show the original contig type as given in CARD v3.1.0 (ncbi_chromosome or ncbi_plasmid). This plot shows only sequences which passed filtering requirements (see Methods). For consistency only sequences between 2000-2022 are shown, omitting a small minority of older sequences (30 for TEM-1, 1 IMP-4). Genera counts are given for common genera, with 26 genera with <50 sequences in the original dataset grouped into ‘Other’ (*Achromobacter*, *Alcaligenes*, *Avibacterium*, *Bacillus*, *Bacteroides*, *Chlamydia*, *Chryseobacterium*, *Cronobacter*, *Haemophilus*, *Leclercia*, *Legionella*, *Morganella*, *Mycoplasma*, *Myroides*, *Neisseria*, *Pasteurella*, *Propionibacterium*, *Providencia*, *Ralstonia*, *Raoultella*, *Serratia*, *Shewanella*, *Shigella*, *Sphingobacterium*, *Stutzerimonas*, *Vibrio*). Full genera and contig counts for each beta-lactamase are provided as Table S1.

Beta-lactamases are a key clinical problem in Gram-negative species (Bush and Bradford, 2020) and recent estimates suggest much higher prevalences of beta-lactamases in the Global South (Antimicrobial Resistance Collaborators, 2022). WHO has designated Gram-negative species priority pathogens for the development of new antibiotics (World Health Organization, 2017). In recent decades, many newly-described beta-lactamases have been identified in clinical bacteria (Bush and Bradford, 2020). A particular concern are emerging extended-spectrum beta-lactamases (ESBLs) which confer resistance to a range of beta-lactams (Livermore, 2008) and commonly spread on MGEs. In some cases these genes can be identified as having been mobilized by MGEs from the chromosomes of environmental bacteria – Partridge (2011) gives a list of the probable ancestral species for many beta-lactamases. Many mobile beta-lactamases are speculated to have originated from a single mobilization event. Mobile beta-lactamases therefore provide repeated examples of a common pattern: mobilization of a chromosomal gene, followed by diversification of its flanking regions as it spreads through bacterial pangenomes into new genomic contexts under strong selective pressure.

We selected twelve clinically important beta-lactamases as focal genes, then downloaded chromosome and plasmid sequences to run through our pipeline looking at +/-5kb flanking regions (see Methods). A single amino acid change in a beta-lactamase is enough to denote a new numbered variant (e.g. *bla*_NDM−1_ and *bla*_NDM−2_) although identical amino acids can still have synonymous SNVs in their nucleotide sequence. We arbitarily chose 25 nucleotide-level differences in the focal gene as a cutoff to include highly-related variants, approximately corresponding to a 97.5% nucleotide identity cutoff (in practice nearly all included had <7 SNVs so >99%, as expected with their recent expansion in human pathogens). Each beta-lactamase gene was seen across multiple genera and all but *bla*_IMP-4_ were found on both chromosomes and plasmids (Fig. 2).

The interactive visualizations produced are available at https://liampshaw.github.io/flanking-regions. For the remainder of this paper, we explore three general quantitative findings that hold – more or less – across these diverse genes, and comment on the possible processes that generate structural diversity.

**FIG. 3.**
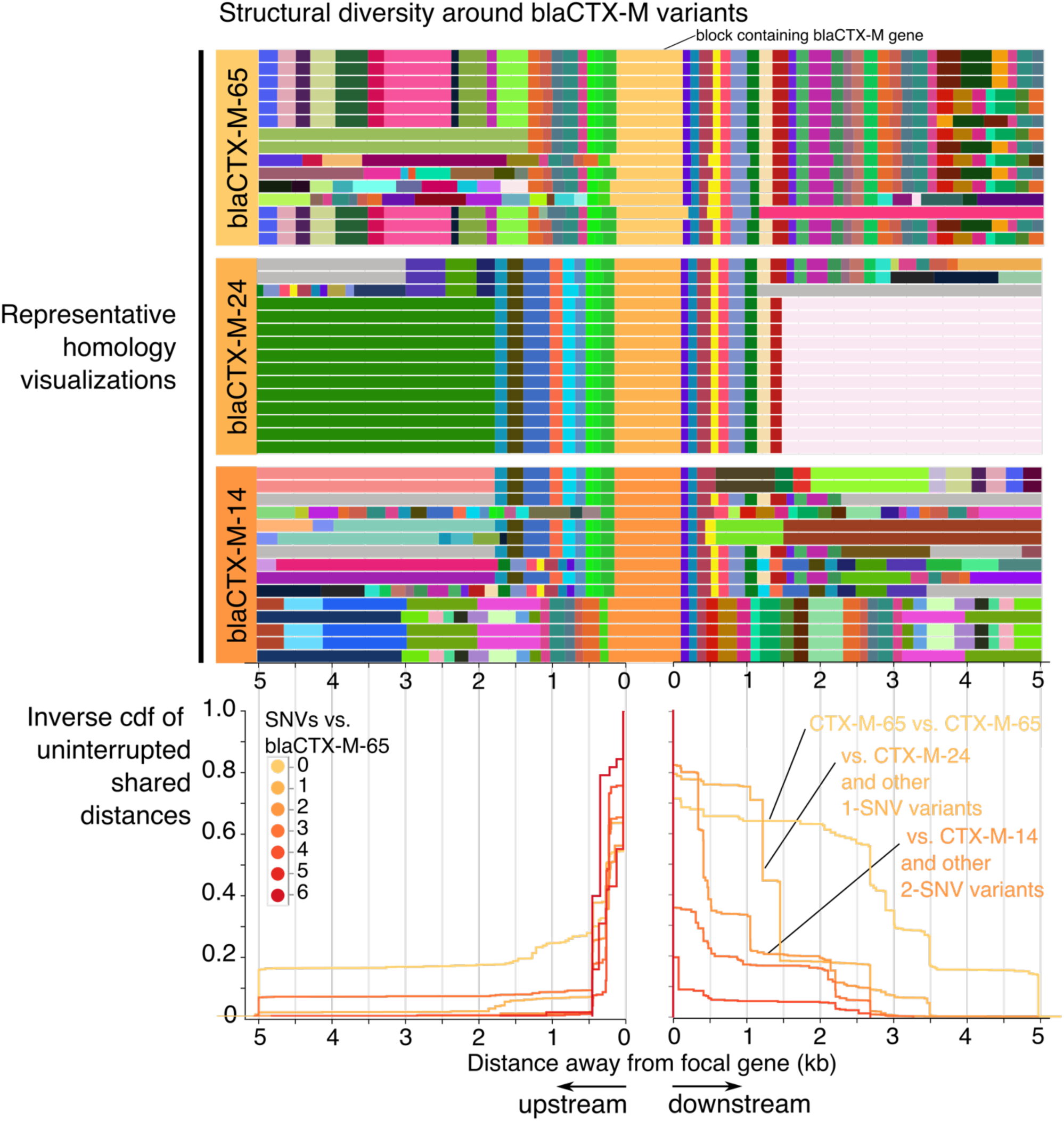
The breakdown of homology in the flanking regions of *bla*_CTX−M−65_ and closely-related genes. The upper panels show the visualization of homology in the 5kb flanking regions of *bla*_CTX−M−65_ and two closely-related genes that encode different CTX-M protein variants, *bla*_CTX−M−24_ (1 SNV apart) and *bla*_CTX−M−14_ (2 SNVs apart). Only a subset of 15 sequences are shown for each gene. Grey blocks have no homology within the wider dataset. Below these visualizations is the inverse cumulative distribution function of pairwise comparisons of distance to the first breakpoint (first occurrence of non-homologous sequence) between *n*=434 sequences carrying any *bla*_CTX−M_ gene with <25 SNVs to *bla*_CTX−M−65_, stratified by the number of SNVs in the focal gene relative to *bla*_CTX−M−65_ (only comparisons involving *bla*_CTX−M−65_ are shown). The same pattern is seen when picking a single isolate for each year/country/genus combination to control for potential sampling bias (Fig. S1).

### 1. The accumulation of nucleotide-level variation correlates with the breakdown of flanking homology

In a simple model of a gene that is mobilized from a chromosomal background and then spreads horizontally on a MGE, the accumulation of single nucleotide variants (SNVs) in the gene itself should be a ‘slow’ molecular clock in contrast to the ‘fast’ rearrangements that happen in its flanking regions. The two would be expected to be broadly correlated: more SNVs in the gene suggest that more time has elapsed, and so greater structural diversity should have accumulated in its flanking regions.

The CTX-M-9-like beta-lactamases provide a clear example of this pattern. Arbitrarily choosing the CTX-M-9-like gene bla_CTX−M−65_ as a focal gene and searching for sequences with variants, we find a strong correlation between SNVs in the gene and uninterrupted shared distances in flanking regions (Fig. 3). This does not appear to be due to repeated sequencing due to outbreaks, because the pattern persists if we include one random isolate from each year/country/genus combination (Fig. S1). There is still some shared sequence around almost all CTX-M-9-like genes, supporting their common origin in some previous mobilisation. Indeed, previously Olson et al. (2005) found a chromosomal beta-lactamase in *Kluyvera georgiana* that shared 100% amino acid identity with CTX-M-14 and was in a 2.7kb region with 99% nucleotide identity to the complex class 1 integron In60, arguing therefore that *K. georgiana* was the likely source for the progenitor of the CTX-M-9-like group through mobilisation. However, since then different CTX-M-9-like beta-lactamases have diverged in their mobilisation.^2^

With a model of a single initial mobilization, all subsequent disruptions of shared sequence are due to subsequent insertions (or deletions) which introduce non-homologous sequence. There are two ‘modes’ of this breakdown of shared sequence in the cumulative distribution of uninterrupted shared distances: gradual decay and sharp breakpoints (Fig. 3). A sharp breakpoint across pairwise comparisons might be suggestive of a consistent ‘common block’, which could be a coherent MGE inserting into multiple genomic backgrounds or a gene cassette in an integron’s gene array; gradual decay is suggestive of a ‘fossilized’ MGE that is undergoing degradation via the introduction of non-homologous sequence in its flanking regions at random points. These patterns may be driven by the same underlying processes. Adjusting the *pangraph* parameters of the pipeline can reduce or increase the sensitivity to non-homologous sequence. Our experience suggests that using the pipeline with default parameters does well at capturing breaks that, when inspected, are clearly caused by ‘large’ insertions/deletions of sequence (i.e. >100bp).

**FIG. 4.**
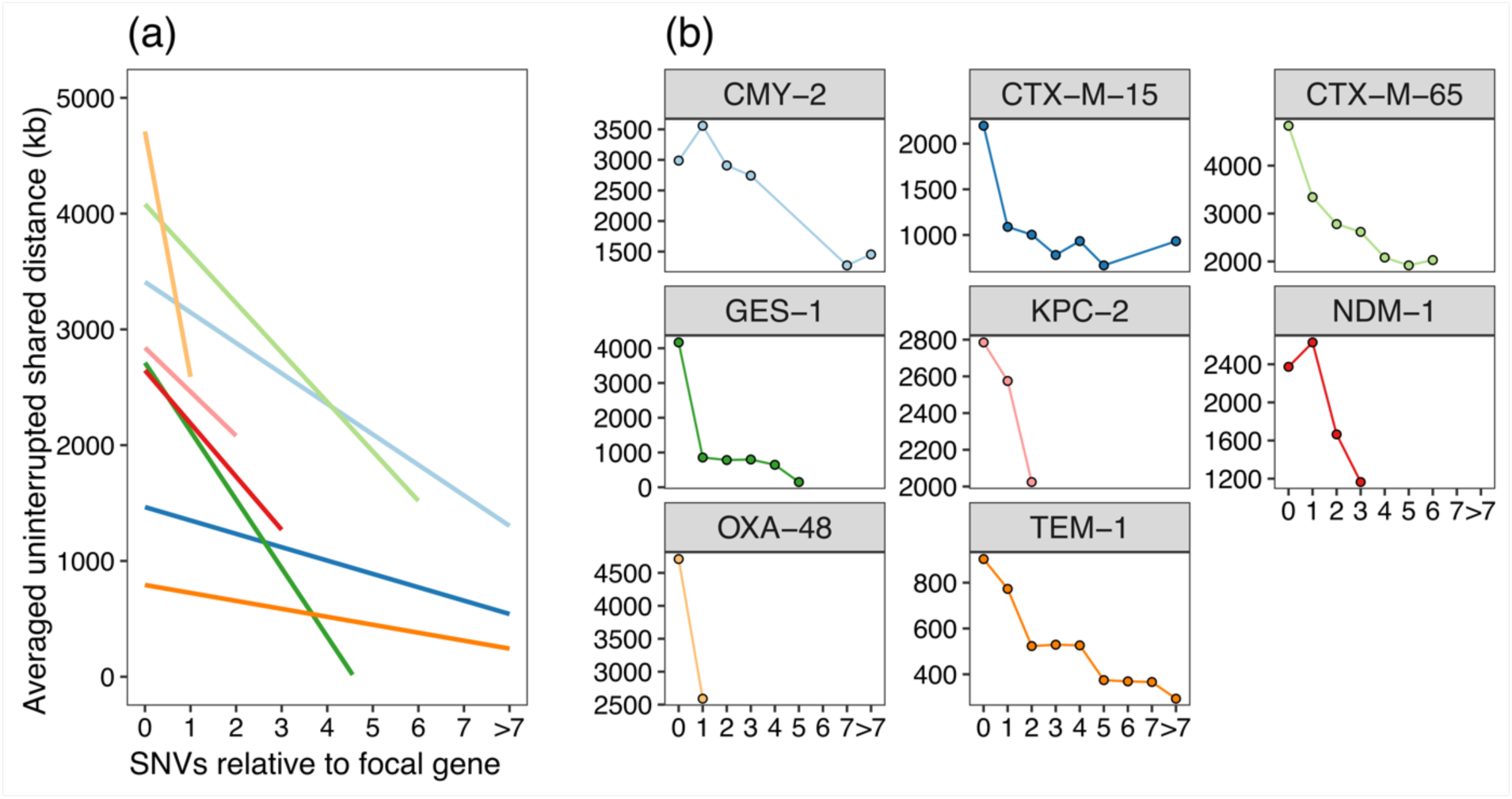
Structural diversity correlates with nucleotide-level variation. Across different betalactamases the presence of any SNV in the focal gene is generally correlated with a shorter flanking region overlap (sum of average upstream and downstream pairwise uninterrupted shared distances). (a) shows linear fits (`stat_smooth`) and (b) average per SNV-level comparison for *n*=7 genes with >100 sequences in the initial dataset (Table I). Not all genes have variants in the dataset for all possible values of SNVs, so where points are absent in (b) this denotes an absence of variants at this difference level. Pairwise comparisons have been deduplicated for year/country/genus combinations before plotting. The flanking region overlap is the sum of the upstream and downstream average uninterrupted shared distances. Number of data points for each gene (after deduplicating): *bla*_CMY−2_ (n=108), *bla*_CTX−M−15_ (n=274), *bla*_CTX−M−65_ (n=119), *bla*_GES−1_ (n=15), *bla*_KPC−2_ (n=161), *bla*_NDM−1_ (n=175), *bla*_OXA−48_ (n=66), *bla*_TEM−1_ (n=514).

The area under the curve (AUC) of the distribution of uninterrupted shared distances for shared sequence between pairs of sequences is equal to the average uninterrupted shared distance length. We can therefore use this to quantify the rough correlation between nucleotide-level variation and structural diversity. By stratifying pairwise comparisons between sequences based on the SNVs in the focal gene, there is a tendency for greater diversification around the focal gene with more SNVs (Fig. 4).

### 2. Linking structural diversity to annotations

It is well-established that insertion sequences (ISs), which use transposases to catalyze DNA cleavage and strand transfer leading to movements, are fundamental to the mobilization of AMR genes and contribute to the complexity of their flanking regions. Repeated ISs are often the reason why flanking regions cannot be assembled in genome assemblies. We used existing annotations from the NCBI gff to identify the locations of ISs as a proxy for IS presence, calculating the transposase density at a position as the number of annotations at that position that contained the term ‘transposase’ divided by the total number of sequences.

Plotting the block diversity and the uninterrupted shared distance distribution with the transposase density revealed interesting patterns (Fig. S2). As an example: for *bla*_NDM−1_, the highest density of transposases is upstream, with an associated immediate breakdown of shared sequence (Fig. 5). After this breakdown of homology, the block diversity is extremely high, indicating that the gene has become stabilized in many different backgrounds. Downstream, the breakdown of the shared ancestral background is more gradual. This recapitulates the more detailed analysis of Acman et al. (2022).

**FIG. 5.**
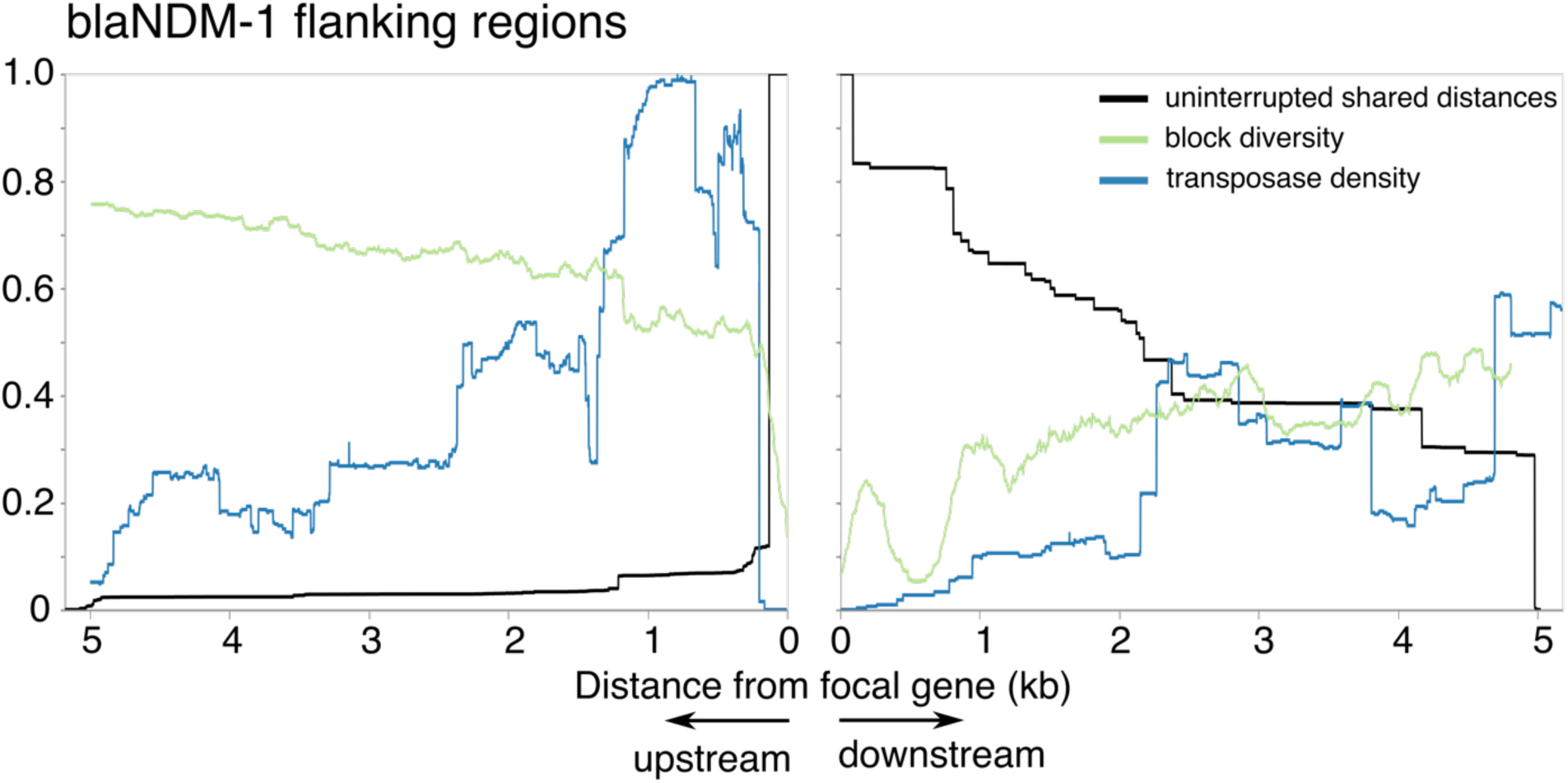
Homology decay, block diversity, and transposase density around *bla*_NDM−1_. Homology decay in terms of pairwise distance to first breakpoint (black), block diversity (green) and transposase density from annotations as a proxy for IS density (blue). All curves are normalized (block diversity to log(N), transposase density to the point of maximum density in the flanking region shown (290/333 sequences). Left shows upstream, right shows downstream.

### 3. A bias for greater structural diversity upstream

Using the distribution of uninterrupted shared distances, we found evidence for an asymmetry between upstream and downstream flanking regions. After removing genes which are known to be found on gene cassettes and associated with integrons (*bla*_GES-1_, *bla*_IMP-4_, *bla*_OXA-10_, and *bla*_VIM-1_; see Partridge et al., 2009), most remaining genes lie below the line of equality, meaning a greater conservation of the downstream flanking region compared to greater structural diversity upstream (Fig. 6). This suggests that for mobile betalactamases associated with ISs, there is a bias for a higher realized rate of introduction of new sequence into the upstream region. Though the numbers are small, it is interesting to note that the exceptions are the oldest genes in the dataset: *bla*_TEM-1_ (first described in 1964) and *bla*_CMY-2_ (1990). The next most recent gene, *bla*_PER-1_ (1993) lies close to the line of equality, with the five remaining genes all described in the 2000s. From this limited evidence, we speculate that this difference between upstream and downstream could be one that is strongest during initial IS-associated spread, fading as genes are recruited into multi-resistance regions.

**FIG. 6.**
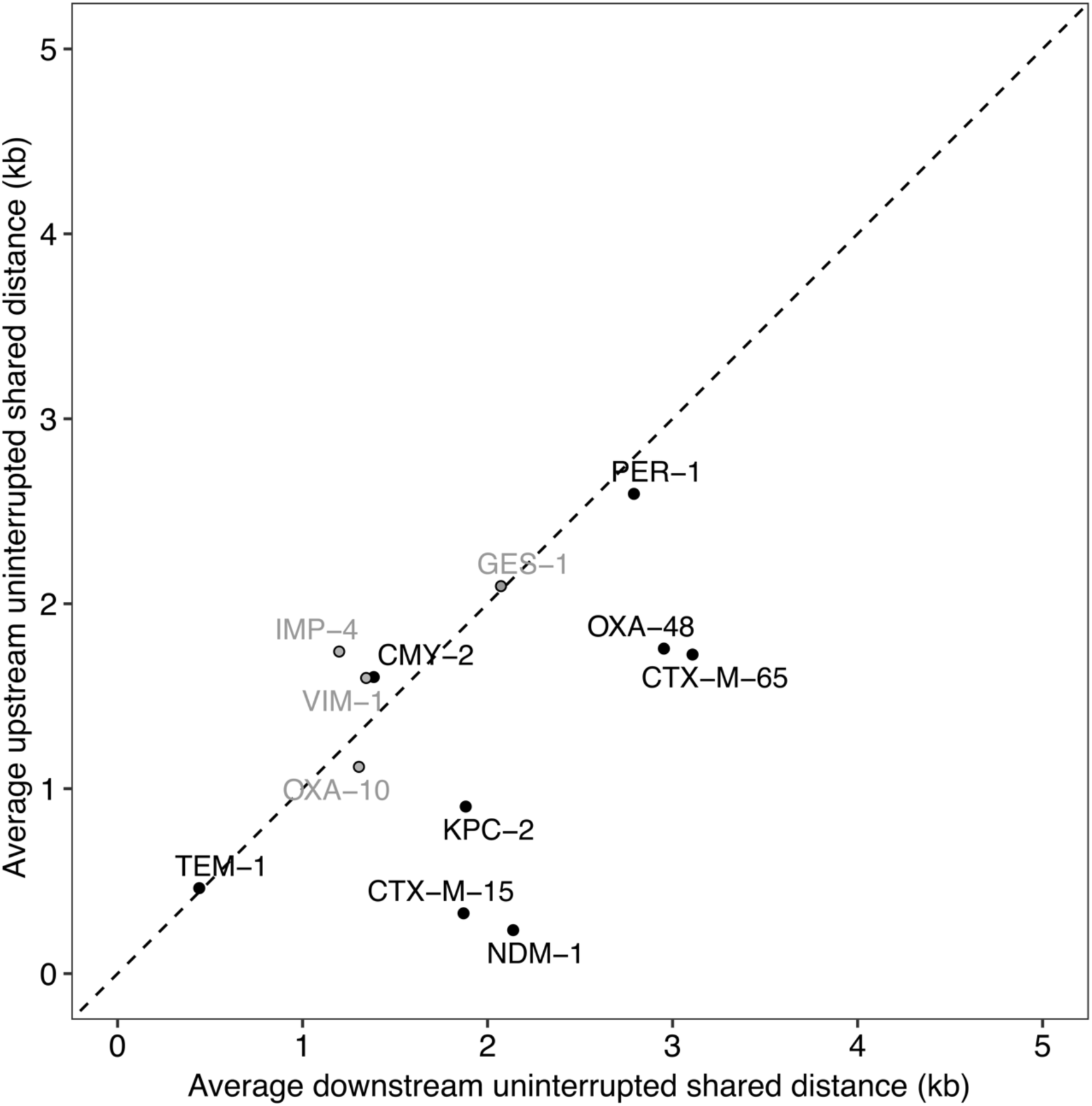
Comparing upstream and downstream uninterruped shared distances. Results show average downstream and upstream pairwise uninterrupted shared distances for each focal gene, including only comparisons between identical focal genes (0 SNVs) after metadata deduplication for year/genus/country. Although most genes lie on the line of equality, there seems to be a bias in some for greater conservation downstream. There is no overall significant difference at α = 0.05 (7/12 downstream>upstream; Wilcoxon signed-rank exact test *V* = 57, *p* = 0.18). However, when integron-associated genes (grey) are removed, the difference becomes stronger (6/8 downstream>upstream, *V* = 32, *p* = 0.055.

## IV. DISCUSSION

We have aimed to provide a starting point to quantitatively investigate structural diversity around mobile genes. By using *pangraph* to find homologous sequences in flanking regions without requiring annotation information, we have constructed a pipeline that quickly produces interactive visualizations that can be explored by researchers to make sense of these complex regions, as well as computing some basic summary statistics. To demonstrate our pipeline, we applied it to the flanking regions of twelve different betalactamase genes in public assemblies. Our observations recapitulate previous knowledge about individual beta-lactamase genes but at scale, and provide evidence of general patterns.

We wish to highlight three limitations of our approach. First, we stress that while this approach gives a way to quantify structural diversity around a gene, it is not intended to infer how a specific instance of the gene is *currently* mobilized. For simplicity we have analyzed 5kb flanking regions, but these are unlikely to capture all information on mobilization: in Gram-negative species mobilized genes tend to cluster in ‘multiresistance regions’ which can be tens of kilobases in size (Partridge et al., 2021). Second, our approach is coarse-grained – in the sense that *pangraph* has a minimum size limit for the size of homologous blocks. This means that traces of evolution at smaller scales will be missed by considering homologous blocks as ‘identical’, for example transposition events which are associated with small-scale (2-bp) insertions. More fine-grained analysis is possible using the multiple-sequence alignments that are generated for each block by *pangraph*, although a bespoke analysis (e.g. an alignment against a known structure to identify small-scale changes) would always be expected to be superior. In particular, an issue in *pangraph* is that breaks in homology involving small blocks close to the minimum size (100bp) appear more unreliable, and we are investigating how best to process these. Third, our analysis uses public genomes which are a biased sample, so all inferences about evolutionary events should be viewed with appropriate scepticism.

We are far from being able to build a quantitative evolutionary model of the structural diversity around mobile genes. However, the patterns we find for beta-lactamases suggest some general principles about the evolution of flanking regions, which appear consistent with what has previously been noted in the literature.

First, nucleotide-level variation in the mobile gene itself does indeed appear to be a ‘slow’ molecular clock compared to the much faster rates of rearrangements in flanking regions. Even single SNVs can change the hydrolytic profile of beta-lactamases, for example in the well-studied evolution of *bla*_TEM-1_ (Barlow and Hall, 2002; Weinreich et al., 2006) including specifically while on MGEs (Rodriguez-Beltran et al., 2018; Kosterlitz et al., 2023), so this variation should not generally be taken as a neutral clock. However, this nevertheless suggests a quantitative connection between mutational and non-mutational evolution that could be fruitful for further exploration. Second, we confirm that ISs can be linked to high levels of structural diversity around mobile genes, likely driven by homologous recombination as well as transposition. Third, we found some genes had an asymmetry with direction, with greater conservation of downstream flanking regions in the majority of non-integron-associated beta-lactamases (6/8), meaning greater structural diversity upstream. Speculatively, we propose that this observation could be a signature of the mobilization dynamics of IS-associated betalactamases.

It is known that many beta-lactamases have been mobilized from chromosomal backgrounds and are weakly expressed in their native genomic context. A wide body of evidence suggests that there is a deep connection between mobility and expression. Not only has antibiotic treatment been shown experimentally to select for the movement of transposon-associated genes from chromosomes to plasmids (Yao et al., 2022), but it was noted as far back as the 1990s that the inverted repeats around ISs often contain promoters, and that such ISs are often found immediately upstream of beta-lactamase genes e.g. for *bla*_TEM−6_ (Goussard et al., 1991) or *bla*_CTX−M−14_ (Cao et al., 2002). Poirel et al. observed that the upstream insertion sequence ISEcp1 which mobilizes *bla*_CTX−M−19_ and other resistance genes contains a strong promoter, simultaneously mobilising the resistance gene and providing strong expression (Poirel et al., 2003). The spread of beta-lactamases is due to strong selection for increased expression: ISs contribute to this in two ways, by providing both a means of transfer onto plasmids and a strong promoter.

Thinking with this simple conceptual model, the initial mobilization of an ancestral beta-lactamase from a chromosome is likely to require the upstream insertion of at least one IS. Therefore, that upstream location will then be a hotspot for the creation of new structural diversity, whether through subsequent insertion to the same target site or homologous recombination between common regions of ISs. We observed that where beta-lactamases are found in gene cassettes and known to be integron-associated (which is the case for *bla*_IMP−4_, *bla*_OXA−10_, *bla*_VIM−1_, *bla*_GES−1_), we did not observe this asymmetry. We also observed that more recent beta-lactamases (described in the 2000s) had more diversity upstream than older genes, suggesting this trend may decay over time as the gene becomes more embroiled in other genetic contexts, with the initial era of spread dominated by a single ancestral MGE coming to an end. Indeed, over time AMR genes tend to accumulate into multi-resistance regions, which has been suggested to be driven by homologous recombination between common components such as transposases (Partridge et al., 2011). Repeated rounds of homologous recombination and insertion that degrade ‘active’ mobile MGEs may play a similar role in the creation of multi-resistance regions to that suggested to be responsible for the creation of defence islands in chromosomes (Rocha and Bikard, 2022).

### Summary

The approach we outline for visualizing and quantifying structural diversity in flanking regions is applicable to any gene and scales to thousands of sequences on the kb-scale. We have applied it to recently emerged beta-lactamases as an example of mobile genes. Each mobile gene is worth detailed study on its own and homology visualizations can help understand the patterns in its flanking regions. Large-scale analysis across genes can reveal patterns consistent with similar underlying processes, suggesting conclusions consistent with the existing literature. Future quantitative methods will allow us to better understand the dynamics that govern the arrival and establishment of genes within pangenomes.

## ACKNOWLEDGEMENTS

The authors thank Sally Partridge, Zamin Iqbal and Will Matlock for helpful feedback on a draft version, and Marco Molari for ongoing and stimulating discussions. Liam Shaw thanks John Lees and Craig MacLean for feedback on visualizations.

## DATA SUMMARY

- Analysis pipeline (release: v1.0.0): https://github.com/liampshaw/mobile-gene-regions. Stably archived at https://doi.org/10.5281/zenodo.10213206
- Interactive plots for n=12 beta-lactamases: https://liampshaw.github.io/flanking-regions
- Beta-lactamase dataset: https://doi.org/10.5281/zenodo.8208376

## Footnotes

1 In line with common usage, by ‘mobile gene’ we refer to any gene that can be or has been found on MGEs in the recent past, on the scale of decades, allowing for the possibility that it may not be *currently* mobile in all genomes.

2 This suggests that *bla*_CTX−M−14_ is the ancestral gene, so would be a better choice of focal gene than *bla*_CTX−M−65_ as we arbitrarily chose here. In fact CTX-M-14 has been suggested to have evolved twice by convergent evolution with different flanking regions for the two nucleotide variants (Navarro et al., 2007).

## REFERENCES

M. Acman, R. Wang, L. van Dorp, L. P. Shaw, Q. Wang, N. Luhmann, Y. Yin, S. Sun, H. Chen, H. Wang, and F. Balloux. Role of mobile genetic elements in the global dissemination of the carbapenem resistance gene *bla*NDM. Nature Communications, 13(1):1131, 2022.

B. P. Alcock, W. Huynh, R. Chalil, K. W. Smith, A. R. Raphenya, M. A. Wlodarski, A. Edalatmand, A. Petkau, S. A. Syed, K. K. Tsang, S. J. C. Baker, M. Dave, M. C. McCarthy, K. M. Mukiri, J. A. Nasir, B. Golbon, H. Imtiaz, X. Jiang, K. Kaur, M. Kwong, Z. C. Liang, K. C. Niu, P. Shan, J. Y. J. Yang, K. L. Gray, G. R. Hoad, B. Jia, T. Bhando, L. A. Carfrae, M. A. Farha, S. French, R. Gordzevich, K. Rachwalski, M. M. Tu, E. Bordeleau, D. Dooley, E. Griffiths, H. L. Zubyk, E. D. Brown, F. Maguire, R. G. Beiko, W. W. L. Hsiao, F. S. L. Brinkman, G. Van Domselaar, and A. G. McArthur. CARD 2023: expanded curation, support for machine learning, and resistome prediction at the Comprehensive Antibiotic Resistance Database. Nucleic Acids Research, 51(D1):D690–D699, 2023.

Antimicrobial Resistance Collaborators. Global burden of bacterial antimicrobial resistance in 2019: a systematic analysis. Lancet, 399(10325):629–655, 2022.

M. Barber. Staphylococcal infection due to penicillin-resistant strains. British Medical Journal, 2(4534): 863–865, 1947.

M. Barlow, B. G. Hall. Predicting evolutionary potential: in vitro evolution accurately reproduces natural evolution of the tem beta-lactamase. Genetics, 160(3): 823–832, 2002.

A. Bauernfeind, R. Jungwirth, S. Schweighart, and M. Theopold. [Antibacterial activity and beta-lactamase stability of eleven oral cephalosporins]. Infection, 18 Suppl 3:S155–167, 1990.

K. Bush and P. A. Bradford. Beta-Lactamase Inhibitors: An Overview. Cold Spring Harb Perspect Med, 6 (8), 2016.

K. Bush and P. A. Bradford. Epidemiology of -Lactamase-Producing Pathogens. Clinical Microbiology Reviews, 33(2):e00047–19, 2020. ISSN 1098-6618. doi:10.1128/CMR.00047-19.

K. Bush and G. A. Jacoby. Updated Functional Classification of beta-Lactamases. Antimicrobial Agents and Chemotherapy, 54(3):969–976, 2010. doi:10.1128/AAC.01009-09.

IV. Cao, T. Lambert, and P. Courvalin. ColE1-like plasmid pIP843 of Klebsiella pneumoniae encoding extended-spectrum beta-lactamase CTX-M-17. Antimicrobial Agents and Chemotherapy, 46(5):1212– 1217, 2002. ISSN 0066-4804. doi:10.1128/AAC.46.5.1212-1217.2002.

N. Datta and P. Kontomichalou. Penicillinase synthesis controlled by infectious R factors in Enterobacteriaceae. Nature, 208(5007):239–241, 1965.

C. L. M. Gilchrist and Y. H. Chooi. clustermap.js: automatic generation of gene cluster comparison figures. Bioinformatics, 37(16):2473–2475, 2021.

S. Goussard, W. Sougakoff, C. Mabilat, A. Bauernfeind, and P. Courvalin. An IS1-like element is responsible for high-level synthesis of extended-spectrum beta-lactamase TEM-6 in Enterobacteriaceae. Journal of General Microbiology, 137(12):2681–2687, 1991. ISSN 0022-1287. doi:10.1099/00221287-137-12-2681.

P. Huovinen, S. Huovinen, and G. A. Jacoby. Sequence of PSE-2 beta-lactamase. Antimicrobial Agents and Chemotherapy, 32(1):134–136, 1988.

L. S. Jones, M. A. Toleman, J. L. Weeks, R. A. Howe, T. R. Walsh, and K. K. Kumarasamy. Plasmid carriage of bla NDM-1 in clinical Acinetobacter baumannii isolates from India. Antimicrobial Agents and Chemotherapy, 58(7):4211–4213, 2014.

A. Karim, L. Poirel, S. Nagarajan, and P. Nordmann. Plasmid-mediated extended-spectrum beta-lactamase (CTX-M-3 like) from India and gene association with insertion sequence ISEcp1. FEMS Microbiology Letters, 201(2):237– 241, 2001.

O. Kosterlitz, N. Grassi, B. Werner, R. S. McGee, E. M. Top, B. Kerr. Evolutionary “crowdsourcing”: alignment of fitness landscapes allows for cross-species adaptation of a horizontally transferred gene. Molecular Biology and Evolution, msad237, 2023.

L. Lauretti, M. L. Riccio, A. Mazzariol, G. Cornaglia, G. Amicosante, R. Fontana, and G. M. Rossolini. Cloning and characterization of blaVIM, a new integron-borne metallo-beta-lactamase gene from a Pseudomonas aeruginosa clinical isolate. Antimicrobial Agents and Chemotherapy, 43(7):1584–1590, 1999.

S. G. Lee, S. H. Jeong, H. Lee, C. K. Kim, Y. Lee, E. Koh, Y. Chong, and K. Lee. Spread of CTX-Mtype extended-spectrum beta-lactamases among bloodstream isolates of Escherichia coli and Klebsiella pneumoniae from a Korean hospital. Diagnostic Microbiology and Infectious Disease, 63(1):76–80, 2009.

D. M. Livermore. Defining an extended-spectrum beta-lactamase. Clinical Microbiology and Infection, 14 Suppl 1:3– 10, 2008.

W. Matlock, S. Lipworth, B. Constantinides, T. E. A. Peto, A. S. Walker, D. Crook, S. Hopkins, L. P. Shaw, and N. Stoesser. Flanker: a tool for comparative genomics of gene flanking regions. Microbial Genomics, 7(9), 2021.

F. Navarro, R. J. Mesa, E. ó, L. mez, B. Mirelis, and P. Coll. Evidence for convergent evolution of CTX-M-14 ESBL in Escherichia coli and its prevalence. FEMS Microbiol Letters, 273(1):120–123, 2007.

N. Noll, M. Molari, L. P. Shaw, and R. A. Neher. : scalable bacterial pan-genome graph construction. Microbial Genomics, 9(6), 2023.

P. Nordmann, E. Ronco, T. Naas, C. Duport, Y. Michel-Briand, and R. Labia. Characterization of a novel extended-spectrum beta-lactamase from Pseudomonas aeruginosa. Antimicrobial Agents and Chemotherapy, 37(5):962– 969, 1993.

A. B. Olson, M. Silverman, D. A. Boyd, A. McGeer, B. M. Willey, V. Pong-Porter, N. Daneman, and M. R. Mulvey. Identification of a progenitor of the CTX-M-9 group of extended-spectrum beta-lactamases from Kluyvera georgiana isolated in Guyana. Antimicrob Agents Chemother, 49(5):2112–2115, 2005.

E. Osano, Y. Arakawa, R. Wacharotayankun, M. Ohta, T. Horii, H. Ito, F. Yoshimura, and N. Kato. Molecular characterization of an enterobacterial metallo beta-lactamase found in a clinical isolate of Serratia marcescens that shows imipenem resistance. Antimicrobial Agents and Chemotherapy, 38(1): 71–78, 1994.

S. R. Partridge, G. Tsafnat, E. Coiera, and J. R. Iredell. Gene cassettes and cassette arrays in mobile resistance integrons. FEMS Microbiological Reviews 33(4):757–784, 2009. doi:10.1111/j.1574-6976.2009.00175.x.

S. R. Partridge. Analysis of antibiotic resistance regions in gram-negative bacteria. FEMS Microbiology Reviews, 35(5), 2011. doi:10.1111/j.1574-6976.2011.00277.x.

S. R. Partridge, Z. Zong, and J. R. Iredell. Recombination in IS26 and Tn2 in the evolution of multiresistance regions carrying blaCTX-M-15 on conjugative IncF plasmids from escherichia coli. Antimicrobial Agents and Chemotherapy, 55(11), 2011. doi:10.1128/AAC.00025-11.

S. R. Partridge, V. I. Enne, E. Grohmann, R. M. Hall, J. I. Rood, P. H. Roy, C. M. Thomas, and N. Firth. Classifying mobile genetic elements and their interactions from sequence data: The importance of existing biological knowledge. PNAS, 118(35):e2104685118, 2021. doi:10.1073/pnas.2104685118.

L. Poirel, I. Le Thomas, T. Naas, A. Karim, and P. Nordmann. Biochemical sequence analyses of GES-1, a novel class A extended-spectrum beta-lactamase, and the class 1 integron In52 from Klebsiella pneumoniae. Antimicrobial Agents and Chemotherapy, 44(3):622–632, 2000.

L. Poirel, C. Heritier, V. Tolun, and P. Nordmann. Emergence of oxacillinase-mediated resistance to imipenem in Klebsiella pneumoniae. Antimicrobial Agents and Chemotherapy, 48(1):15–22, 2004.

L. Poirel, J.-W. Decousser, and P, Nordmann. Insertion Sequence ISEcp1B Is Involved in Expression and Mobilization of a blaCTX-M beta-Lactamase Gene. Antimicrobial Agents and Chemotherapy, 47(9):2938–2945, 2003. ISSN 0066-4804. doi:10.1128/AAC.47.9.2938-2945.2003.

E. P. C. Rocha and D. Bikard. Microbial defenses against mobile genetic elements and viruses: Who defends whom from what? PLOS Biology, 20(1):1–18, 2022. doi:10.1371/journal.pbio.3001514.

J. Rodriguez-Beltran, J. C. R. Hernandez-Beltran, J. DelaFuente, J. A. Escudero, A. Fuentes-Hernandez, R. C. MacLean, R. Peña-Miller, A. San Millan. Multicopy plasmids allow bacteria to escape from fitness trade-offs during evolutionary innovation. Nature Ecology & Evolution 2:873–881, 2018.

A. E. Sheppard, N. Stoesser, I. German-Mesner, K. Vegesana, A. Sarah Walker, D. W. Crook, and A. J. Mathers. TETyper: a bioinformatic pipeline for classifying variation and genetic contexts of transposable elements from short-read whole-genome sequencing data. Microbial Genomics, 4(12), 2018.

M. A. Toleman, J. Spencer, L. Jones, and T. R. Walsh. blaNDM-1 is a chimera likely constructed in Acinetobacter baumannii. Antimicrobial Agents and Chemotherapy, 56(5):2773–2776, 2012.

D. M. Weinreich, N. F. Delaney, M. A. Depristo, D. L. Hartl. Darwinian evolution can follow only very few mutational paths to fitter proteins. Science, 7;312(5770):111–4, 2006.

World Health Organization. Prioritization of pathogens to guide discovery, research and development of new antibiotics for drug-resistant bacterial infections, including tuberculosis (who/emp/iau/2017.12). 2017.

Y. Yao, R. Maddamsetti, A. Weiss, Y. Ha, T. Wang, S. Wang, L. You. Intra- and interpopulation transposition of mobile genetic elements driven by antibiotic selection. Nature Ecology and Evolution. 6(5):555–564, 2022.

H. Yigit, A. M. Queenan, G. J. Anderson, A. Domenech-Sanchez, J. W. Biddle, C. D. Steward, S. Alberti, K. Bush, and F. C. Tenover. Novel carbapenem-hydrolyzing beta-lactamase, KPC-1, from a carbapenemresistant strain of Klebsiella pneumoniae. Antimicrobial Agents and Chemotherapy, 45(4):1151–1161, 2001.

D. Yong, M. A. Toleman, C. G. Giske, H. S. Cho, K. Sundman, K. Lee, and T. R. Walsh. Characterization of a new metallo-beta-lactamase gene, bla(NDM-1), and a novel erythromycin esterase gene carried on a unique genetic structure in Klebsiella pneumoniae sequence type 14 from India. Antimicrobial Agents and Chemotherapy, 53(12):5046–5054, 2009a.

D. Yong, M. A. Toleman, C. G. Giske, H. S. Cho, K. Sundman, K. Lee, and T. R. Walsh. Characterization of a new metallo-beta-lactamase gene, bla(NDM-1), and a novel erythromycin esterase gene carried on a unique genetic structure in Klebsiella pneumoniae sequence type 14 from India. Antimicrobial Agents and Chemotherapy, 53(12):5046–5054, 2009b.

