## Supplementary Material for "Visualizing and quantifying structural diversity around mobile resistance genes"

Liam P. Shaw[1, 2, *] and Richard A. Neher[3]
[1] *Department of Biology, University of Oxford, Oxford, UK*
[2] *Department of Biosciences, University of Durham, Durham, UK*
[3] *Biozentrum, University of Basel, Basel, Switzerland*

**Table S1.** Summary of the genomes included in the filtered dataset for each of the twelve beta-lactamases, giving counts for different bacterial genera over time and the numbers of chromosomes/plasmids.

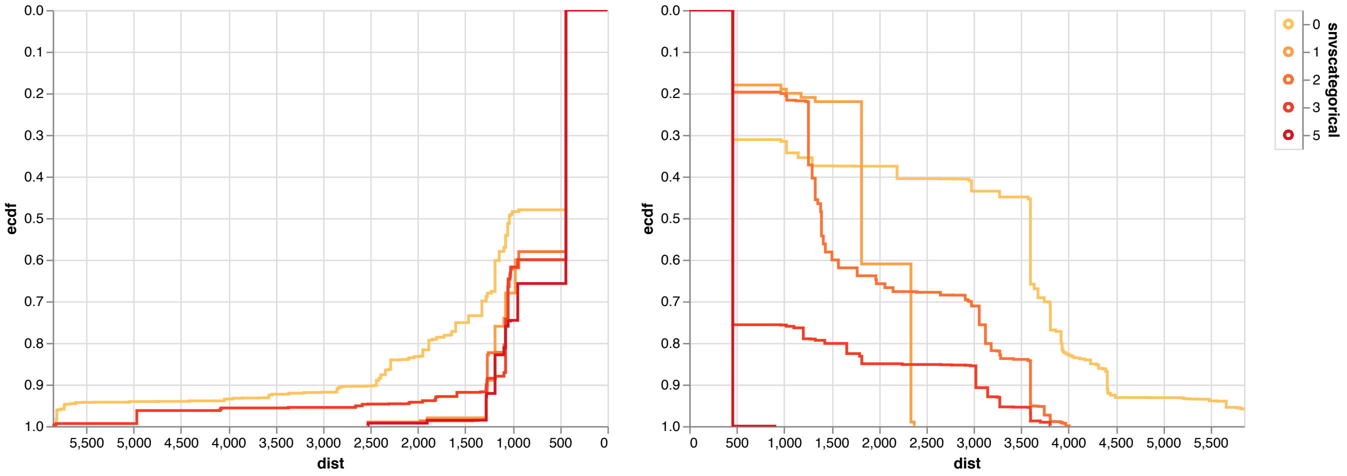


FIG. S1: **The breakdown of homology in the flanking regions of *bla*$_{\mathrm{CTX-M-65}}$ and related genes.** As in the main manuscript Fig. 2b, but here we pick only a single isolate for each year/country/genus combination to control for potential sampling bias.

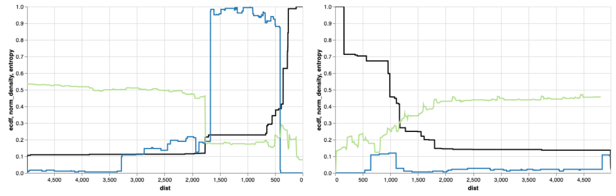

(a) blaCMY-2

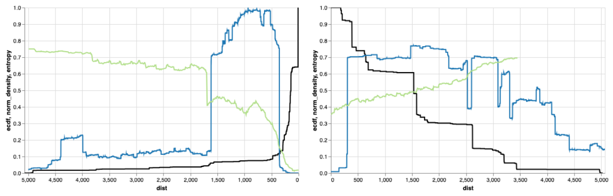

(b) blaCTX-M-15

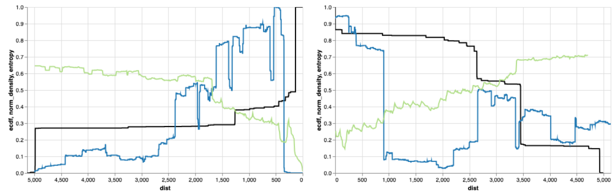

(c) blaCTX-M-65

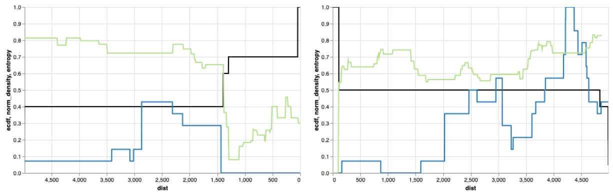

(d) blaGES-1

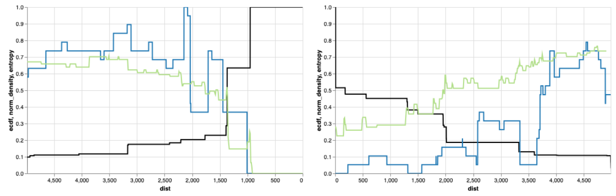

(e) blaIMP-4

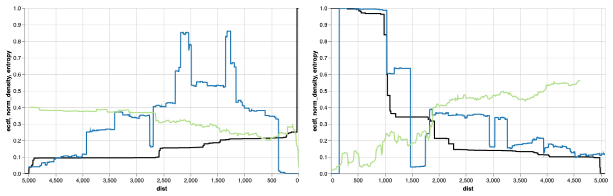

(f) blaKPC-2

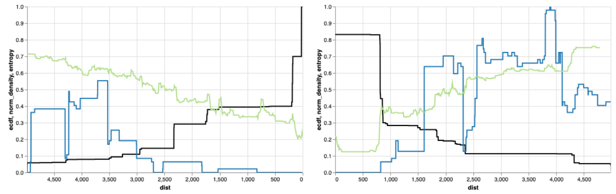

(g) blaOXA-10

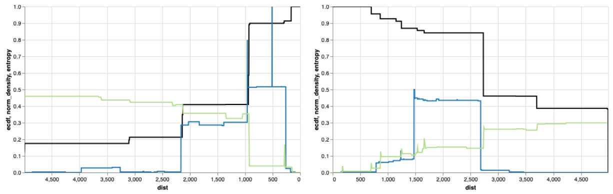

(h) blaOXA-48

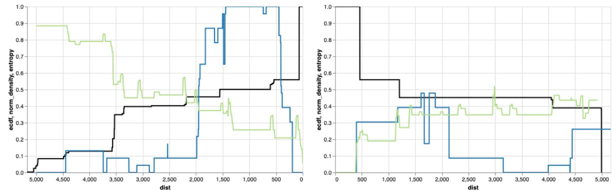

(i) blaPER-1

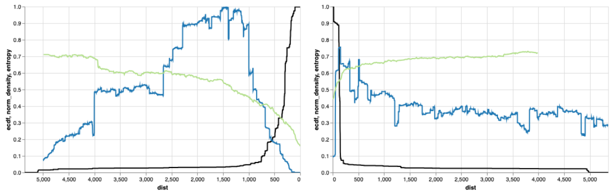

(j) blaTEM-1

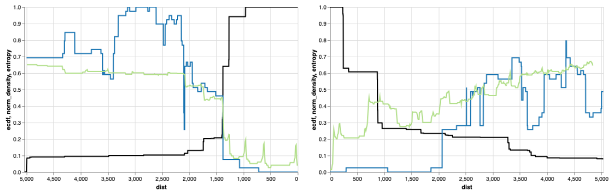

(k) blaVIM-1

FIG. S2: **Flanking regions of eleven beta-lactamase genes.** Similar to the plot for blaNDM-1 in the main text: overlaid plots of the inverse cdf of uninterrupted shared distances (black), with normalised block diversity (green) and transposase density (blue) as in the main text.
